# Statistical power of gene-set enrichment analysis is a function of gene set correlation structure

**DOI:** 10.1101/186288

**Authors:** David M. Swanson

**Author notes:** To whom correspondence should be addressed. ORCID ID: 0000-0003-3174-1656.

## Abstract

**Motivation:** We describe why statistical power for both self-contained and competitive gene-set tests is a function of the correlation structure of co-expressed genes, and why this characteristic is undesirable for gene-set analyses. Variable statistical power as a function of gene correlation structure has not been observed or studied previously. The observation is important in part because gene-set testing methodology is well-developed, yet this fundamental feature of many of its tests is unknown and has the potential to reinterpret past gene-set test results and guide future implementations, including those using sequence data. Type 1 error inflation is also amenable for study in our statistical framework; while it has been well-studied and described previously for both self-contained and competitive tests, it has less often been done in an analytical framework. Our observations apply to four commonly-used gene-set testing approaches for microarrays, including CAMERA, ROAST, SAFE, and GAGE, and a recently proposed one for RNAseq, MAST.

**Results:** We characterize situations in which power is especially small relative to effect sizes of genes in a set for both competitive and self-contained gene-set tests. We propose three alternative tests, one of which replicates the properties of permutation-based self-contained tests, but avoids the need for even recently proposed, rotation-based approximations to permutations. The two other proposed tests have the unique property that statistical power is not a function of co-expression correlation in the gene-set and therefore is the preferred methodology. We compare our proposed tests to leading gene-set tests and apply them to an already-published study of smoking exposure on pregnant women.

**Supplementary Material:** Online supplementary material includes additional simulation results supporting the relationship between the “mixed” and “directional” gene-set tests of ROAST and closed-form implementations of them.

## 1 Introduction

Gene set enrichment analysis (GSEA) or gene-set testing is a class of statistical methods whose goal is to assess the joint enrichment of a biologically interpretable set of genes in a microarray or RNAseq experiment^1–6^. A variety of methods have been proposed nearly since the advent of microarrays, all with unique advantages, and there have been more recent modifications of these methods as RNAseq becomes a more common platform^7^. There have been two primary categories of gene-set tests, “competitive” and “self-contained” ^8–11^. The former tries to answer whether genes in a set are more differentially expressed than some background level of differential expression on the array. The latter tries to answer whether the gene-set is more differentially expressed than one were to expect under the null of no association between transcript abundance and experimental condition. The null distributions for these two kinds of hypothesis tests have often been calculated via permutation, either at the gene-level for the competitive test, or sample level, for the self-contained test. Manual permutation is used in an attempt to attain the nominal type 1 error rate^12,13^ or sometimes computationally easier methods that yield similar results^14,10^. Rejecting a competitive gene-set test test is often a higher bar or more difficult burden of proof, and these tests have been more commonly used in gene-set testing. A rejection of a competitive gene-set test can be more biologically meaningful than a self-contained test.

In this paper, we describe within an analytic statistical framework why past gene-set tests have suffered from inflated type 1 error, and how permutation-based methods have sought to address the issue in different ways. Goeman and Buhlmann ^9^ made a similar and rigorous description of gene-set tests, though in the context primarily of 2×2 tables, and prior to the development of some of the tests we analyze. More importantly, we additionally describe in the same framework why past and more recently proposed gene-set tests suffer from having variable power as a function of gene set correlation structure.

We show that the location of causal transcripts within the co-expression correlation structure is critical in determining statistical power of the test in addition to the effect sizes of them, and how the issue is relevant to both competitive and self-contained gene-set tests. We define causal transcripts as those expressed genes having a causal effect on the outcome of interest, rather than only being correlated with those that are. Briefly, correlated blocks of genes manifest themselves in disproportionate degree in the tails of distributions under both the null and alternative hypotheses. Since we generally perform statistical tests in the tails of distributions, hypothesis tests are disproportionately influenced by this correlation. When causal transcripts are found in these correlated blocks, power for their detection is likewise overly represented relative to causal transcripts found in less correlated blocks. This observation has been made in the context of region-based SNP testing, though not in the context of gene-set testing^15^. Despite the observation being made in the context of SNP-set testing, it is pertinent to mention since tests continue to be proposed that suffer from variable power under different correlation structures^16^.

We leverage our statistical analysis of gene-set tests by proposing three tests, the suite of which we call “JAGST” (Just Another Gene Set Test). The first test is self-contained and replicates results from a leading self-contained test ROAST, but does so using a closed form mixture density rather than the rotation and iteration-based method of ROAST. Though ROAST is an elegant, general, and computationally cheap algorithm for various set-based hypothesis tests, two of its more important special cases (mixed and directed set-based tests) fall within our statistical framework so that relevant null tail probabilities can be written down and calculated.

The second two tests of JAGST are power-invariant self-contained and competitive tests. The proposed JAGST power-invariant self-contained test simply uses correlation information of differential expression test statistics to implement a standard chi-square test. The proposed competitive test is more complex and computationally intensive, using a sophisticated and recently proposed high-dimensional penalized regression method for obtaining z-statistics from a variable selection procedure ^17–20^. We use these z-statistics to create an appropriate mixture distribution of non-central chi-squares, which is the null distribution for our competitive test.

We perform simulations with leading and commonly used competitive and self-contained tests, including ROAST, CAMERA, SAFE, and GAGE^8,14,13,6^ to demonstrate the degree to which statistical power is effected by correlation structure of the gene-set and how our proposed tests do not have this same property. ROAST and SAFE are self-contained tests, while CAMERA and GAGE are competitive tests. We perform a data analysis on an already published study of smoking exposure in pregnant women ^21^. While expression studies are the context of our analysis, our observations are also relevant to RNAseq. The popular MAST method for RNAseq is based on CAMERA, and many of the same statistical principles will apply in the RNAseq context^7^. Additionally, expression analyses continue to be a valuable and well-used platform for diagnosis and prediction^22–26^.

## 2 Methods

Consider matrix Y (*n* × *m*) of normalized transcript measures and matrix X (*n* × *p*) of experimental conditions, where *Y* has *m* × *m* correlation matrix Σ, whose *i, j*^*th*^ element is *ρ*_*i.j*_. Assume for now that *p* = 1 for simplicity. It is common in GSEA to first perform a standard microarray analysis, and then test gene-set enrichment on the summary statistics (such as transcript-level test statistics).

If we fit a regression model of the *i*^*th*^ column of Y (*Y*_*i*_) on X, *i ∈* 1 *… m*, we extract some test statistic, call it *T*_*i*_ (oftentimes a t-statistic for linear regression if the outcome is continuous), associated with this model, and likewise *T*_*j*_ for the model of *Y*_*j*_ on X. The test statistic from a linear regression model is

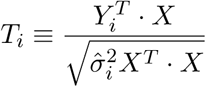

if we have centered *X* and *Y*, which follows a *t*-distribution on *n*-2 degrees of freedom under the null hypothesis and where 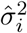 is the estimated variance of *Y*_*i*_ conditional on *X*. Likewise, define

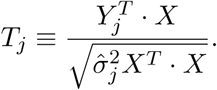

Under the null hypothesis of no association between *Y*_*i*_ or *Y*_*j*_ and *X, E*[*T*_*i*_] = *E*[*T*_*j*_] = 0. Since *Cor*(·, ·) is invariant to scaling its arguments and assuming centered, normalized *Y*_*i*_’s, *Y*_*j*_’s, and *X*’s,

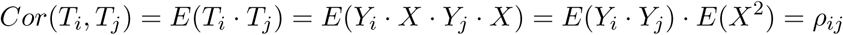

where the second to last equality holds because we calculate under the null hypothesis of no association with *X*. So we see that *Cor*(*T*_*i*_, *T*_*j*_) is the (*i, j*) entry of Σ, the correlation matrix of *Y*. It follows that the *m-*length vector *T* composed of entries *T*_*k*_, *k* = 1, *…, m* also has *Cor*(*T*) = Σ.

Now consider the regression model under the alternative hypothesis of an association between an experimental condition and transcript abundance. In particular, suppose there exists a causal association between *X* and *Y*_*j*_ such that the expectation of the test statistics *T*_*j*_ corresponding to regression model *Y*_*j*_ ∼ *X* is *E*[*T*_*j*_] = *μ*_*j*_. For example, power under this association for an *α*-level test would be calculated with

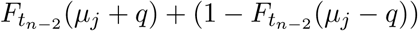

where 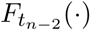 is the cdf of a t-distribution on *n*-2 degrees of freedom and *q* is the *α/*2 quantile of that distribution. The expression is the sum of two tail probability regions which corresponds to the rejection region in a two-sided test and is calculated under the alternative hypothesis of *E*[*T*_*j*_] = *μ*_*j*_.

Assume there exists no causal association between *X* and *Y*_*i*_, and consider test statistic *T*_*i*_. Because *Cor*(*T*_*i*_, *T*_*j*_) = *ρ*_*i,j*_, *E*(*T*_*i*_) = *ρ*_*i,j*_ *· μ*_*j*_; we have power to detect an association between *X* and *Y*_*i*_ by virtue of *Y*_*i*_’s correlation with *Y*_*j*_. though less so because *|ρ*_*i,j*_*|* ≤ 1. We will return to this point later in the discussion of statistical power.

Suppose that we have sufficient sample size such that the t-statistics resulting from an expression analysis can be approximated by the standard normal distribution. Since the test statistic for many gene-set tests relies on or is highly correlated with a sum or sum of squares of probe-level summary statistics composing the gene-set (e.g,^14,3,8,27,6,13^), consider the following property about the multivariate normal distribution.

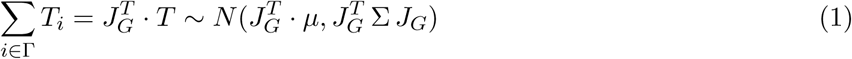

for *T*, as above, the length *m* vector that follows *MV N* (*μ*, Σ), Γ a size *g* set of indices for the gene-set, and *J*_*G*_ a length *m* indicator vector whose 1 entries correspond to the indices of Γ. So the mean of Σ_*i∈*Γ_ *T*_*i*_ is the sum of the component means corresponding to the 1 entries in *J*_*G*_, and the variance of Σ_*i∈*Γ_ *T*_*i*_ is the sum of the entries of Σ that remain after multiplication by *J*_*G*_. Sometimes this test statistic in the gene-set testing context is called the directed or directional test since its value is sensitive to the sign of *T*_*i*_.

Additionally, if we square the *T*_*i*_’s, we have

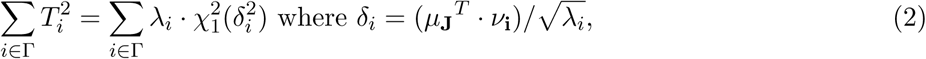

*μ*_*J*_ *= J*_*G*_ *·μ*, and *λ*_*i*_ and *ν***_i_** are the eigenvalues and eigenvectors of 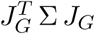^28–30^. Sometimes this test statistic in the gene-set testing context is called undirected, non-directional, or mixed as its power is invariant to the sign of *T*_*i*_.

## 3 Relationship to permutation-based tests

### 3.1 Considering preservation of Type 1 error using permutations

It is sometimes thought that permutation is a sure, if computationally burdensome, way of preserving type 1 error when the null distribution of a test statistic is unknown. We show why this is not the case, at least with competitive gene-set tests. The result has been noted frequently elsewhere^8,27,10^, but we try to describe it in a statistical framework so that evidence of the phenomenon is not primarily simulation-based. First, however, we consider permutation-based self-contained tests and the parametric distributions approximated by them.

#### 3.1.1 Self-contained tests type 1 error

In light of the results in Section 2, we can consider the null distributions generated when samples (“self-contained” gene-set tests) or genes (“competitive” gene-set tests) are permuted to generate a null distribution for the test statistics discussed. In permuting samples, any possible association between gene and outcome is broken so that the expectation of *T*, the test statistic vector described above, is a length *m* vector with expectation **0**, rather than *μ*. Since the permutations do not permute the particular gene set under consideration, the correlation structure of the gene-set is preserved. Therefore if we define 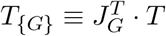, one can conclude that a permutation-generated null distribution for *T*_{*G*}_ is a sample from

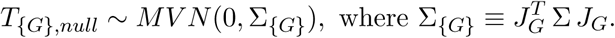

So Σ_{*G*}_ is the correlation matrix of the transcripts, *Y*_*i*_’s, in the gene-set. With the joint distribution of *T*_{*G*},*null*_ in mind, we can consider the sum of squares of its elements. Since the expectation of the vector *T*_{*G*},*null*_ is zero and using notation conventions from Section 2, *δ*_*i*_ = 0 because *ν*_*i*_ will be an inner product with a **0** vector. Differences in the null distribution for 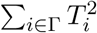 are therefore driven by {*λ*_*i*_} (the set of eigenvalues of Σ_{*G*}_), and the null distribution can be very different depending on the distribution of the {*λ*_*i*_}’s. If there is high within-set correlation, greater variation in *λ*_*i*_ will lead to a heavier-tailed null distribution, whereas little within-set correlation will correspond to relatively similar *λ*_*i*_’s and a thinner tail.

No within gene-set correlation implies independent *T*_*i*_’s, so that

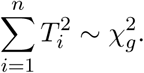

It is because independent component test statistics have been assumed at times in the past that a 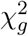 null distribution is used when in fact the heavier tailed 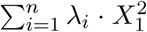 was the true null distribution^4^. It is in this distinct way that even self-contained tests have occasionally had inflated type 1 error, though this has been recognized and corrected in different articles^31,14,10,32^.

#### 3.1.2 Competitive tests type 1 error

When permuting across gene sets for a competitive gene set test, probe-wise associations with the outcome are maintained, but the set over which we aggregate changes as a function of the genes randomly chosen to compose our permutation-generated set. Thus, for permutation *k* of the null distribution generation, the distribution from which we sample is

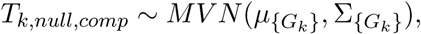

where {*G*_*k*_} is a set of genes of size *g* randomly chosen, 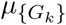 is the expectation vector of test statistics corresponding to those particular genes, and 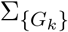 is a submatrix of Σ corresponding to the gene set {*G*_*k*_}. Since {*G*_*k*_} changes with each iteration, it is evident that {*T*_1,*null,comp*_, *…, T*_*q,null,comp*_}, where *q* is the size of the permutation-generated null, are not identically distributed, but a mixture of distributions.

Crucially, and likely the reason type 1 error has been difficult to maintain in competitive gene-set tests, the off-diagonal entries of 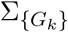 (the correlation matrix of permutation *k*) will often be systematically smaller than the off-diagonal entries of 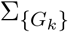 because genes within biologically meaningful sets will often be co-expressed to a greater degree than randomly chosen groups of genes. We may therefore expect that the permutation-generated null distribution will not be as thick tailed as the test statistic whose null it is trying to approximate.

### 3.2 Statistical power when using permutations

Because expression analyses often rely on univariate regression test statistics, confounding structures are not detected and power to detect some non-causal transcripts can be above *α* because of their association with the causal gene. The underlying causal structure of transcript on phenotype and confounding by co-expressed genes and their effects on statistical power has been less discussed on a methodological level. When variable power has been discussed, it has been done so from the perspective of underlying biological or platform phenomena, such as in Oshlack and Wakefield ^33^ with respect to transcript length bias. Here we describe why statistical power for both self-contained and competitive gene-set tests can vary due to the methodological set-up of the tests themselves.

#### 3.2.1 Illustrative example

We first consider an example to explain variable power more concretely. For simplicity, assume a gene-set consists of 3 genes, one of which has a causal association with the outcome, and the other two do not. Call the test statistic associated with the causal gene *T*_1_ with expectation *μ*_1_, and those of the non-causal genes *T*_2_ and *T*_3_ (both with expectations 0). First suppose that *T*_2_ and *T*_3_ have correlation *ρ*, but both are independent of *T*_1_ (the causal gene). The distribution of their sum is

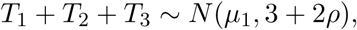

while the non-directional test statistic is

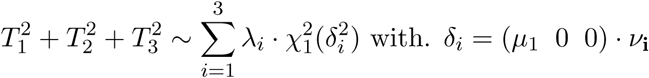

and *λ*_*i*_ and *ν***_i_** are the eigenvalues and eigenvectors of Σ.

Consider now the scenario where we “shift” the correlation structure so that *T*_1_ and *T*_2_ have correlation *ρ*, and *T*_3_ is independent of them both. though the same causal associations remain. So *E*(*T*_1_) = *μ*_1_, and in this scenario *E*(*T*_2_) = *μ*_1_*ρ* because of *T*_2_’s correlation with *T*_1_. The expectation of *T*_3_ is still 0. This time, the distribution of their sum is

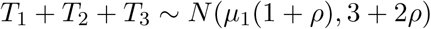

and the non-directional test statistic is

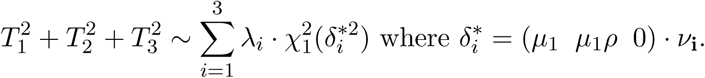

It will always be the case that 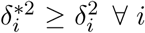, that is, in the second scenario the non-centrality parameters will tend to be bigger than those in the first scenario. We know this because (*μ*_1_ *μ*_1*ρ*_ 0) is greater than (*μ*_1_ 0 0) element-wise. Additionally, there will exist at least one *δ*_*i*_ strictly greater in the second scenario than the first scenario because for at least one *i, ν*_*i*_ has a non-zero entry in the second element since eigenvectors span the space defined by the columns of Σ. Because *χ*^2^ distributions are stochastically strictly increasing in their non-centrality parameters, we will always be better powered to detect the gene-set in the second scenario than in the first.

From this example, we conclude that we are more likely to reject the null for gene set enrichment test when causal transcripts are co-expressed with non-causal ones.

## 4 Proposed hypothesis tests: JAGST

### 4.1 Proposed hypothesis test 1: an analytic approach to ROAST’s self-contained test

First we introduce directed and mixed versions of self-contained gene-set test that are nearly numerically equivalent to these important special cases of ROAST’s versions of them^14^. Our directed and mixed tests rely on closed form formulae rather than the rotation-based methodology of ROAST, however. ROAST is itself an elegant way of approximating the results of permutation-based procedures.

The test statistics for the directed and mixed self-contained gene-set tests were in fact already introduced in Section 2 with equations (1) and (2), respectively. Equation (1) is the sum of differential expression test statistics, while equation (2) is the sum of squares of differential expression test statistics. Since the null distributions of (1) and (2) are known and provided in the expressions, we do not need to rely on the rotation of residual methodology, which though statistically elegant has p-value granularity dependent on the number of specified iterations. We can instead calculate p-values using tail probabilities of the normal distribution in the case of the directional test (eqn (1)) and of a mixture of scaled *χ*^2^ distributions in the case of the mixed test (eqn (2)).

We see in the Supplementary Material the correlation of 0.99 (0.96) between the ROAST directional test (mixed test) and analytic calculation of tail probabilities using these densities, both under the null hypothesis. That correlation falls slightly under the alternative hypothesis for reasons explained in the figures, but remains above 0.88. If the variable power issue is not a concern for the analyst, for example for reasons given in the Discussion section, and p-values from the self-contained test ROAST methodology are preferred, one need use only the test statistics and explicit null distributions in (1) or (2). Since inversion of the *χ*^2^ mixture distribution can be difficult, sampling from the mixture distribution is also adequate and likely still more computationally inexpensive than ROAST.

### 4.2 Proposed hypothesis test 2: power-invariant competitive gene-set test

We propose the JAGST competitive test whose statistical power is invariant to the correlation structure in which causal effects are found. We do so by making two key changes to currently used gene-set testing methods. First, define *T*_{*G*}_ and Σ_{*G*}_ as the vector of differential expression test statistics corresponding to gene set {*G*}, and the correlation matrix of the *Y*_*i*_’s making up the gene-set, respectively. Then the test statistic for the competitive test is calculated using

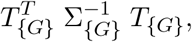

rather than a straightforward sum or sum of squares of test statistics. Using the inverse covariance matrix of the test statistics is central to achieving the desired correlation structure power invariance. Secondly, the null distribution is calculated by taking random subsets of genes of size g and calculating the same test statistic. Unlike other competitive tests that calculate their null distribution with random subsets of genes, using the inverse correlation matrix makes each realization of our null distribution a function of only the underlying effects of each subset thereby controlling for correlation structure.

While this is the essential idea of the JAGST competitive test, in practice we generate the proposed null distribution in a different and computationally easier way. Especially for large gene-sets, some of which approach a few hundred genes, calculating the null distribution in this brute force way is too cumbersome if not numerically prohibitive depending on whether the correlation matrices are approximately singular.

We therefore propose instead to:

1. Perform variable selection on the random gene subsets of size g,
2. choose the model with the smallest AIC (or another model fitness criterion that could be shown to control type 1 error and be power-invariant), then
3. use a novel and sophisticated method to calculate the z-statistics of the selected variables^19,18,20^, in order to
4. take the sum of squares of these test statistics, calling them 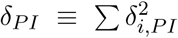 (“PI” for “power-invariant”), and
5. sample from 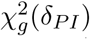, which is the null distribution of the above test statistic as demonstrated in simulation (Figure 4).

We iterate until desired granularity in the null distribution is achieved.

The L1-penalized regression hypothesis testing procedure of Taylor and Tibshirani ^19^ proves central to our algorithm. It provides a means of obtaining z-statistics on selected variables in a penalized regression framework and until recently was not possible. These z-statistics allow us to determine the distribution under the alternative hypothesis that our differential expression test statistics arise from.

While variable selection is not without computation cost, since the procedure stops once a certain AIC is achieved, we will generally avoid the large O(*g*^3^) cost of matrix inversion and numeric instability for large gene sets by a significant margin. This is especially true if it is assumed that in general gene sets are composed of many transcripts working together in pathways, though reveal relatively sparse models when regularization is applied.

Iteration over different sets of size *g* will yield a mixture of non-central *χ*^2^ distributions. Formulae and approximations have been proposed for mixtures of *χ*^2^ distributions^29,34^, which could be used if the same null distribution were relevant to many different tests, or sampling from distributions were burdensome. For our purposes, we eschew these formulae to prefer sampling from the mixture distribution in R^35^.

### 4.3 Proposed hypothesis test 3: power-invariant self-contained gene-set test

For the JAGST self-contained test, we again use the test statistic

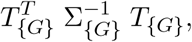

with notation used as before. In this case however, the null hypothesis assumes no association between transcript and outcome so the null distribution for this test statistic is simply 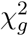. The quantiles for this null distribution will often be a much lower threshold for statistical significance as compared to the proposed competitive gene set test.

## 5 Results

We provide simulation and data analysis results for the different JAGST tests and compare them with CAMERA (with and without ranks), GAGE (with and without ranks), ROAST, and SAFE. We abbreviate our simulation analysis of type 1 error since it has been described well and studied in detail elsewhere^13,36,10,8^ and focus on power as a function of correlation structure under different generating models. We perform simulation and data analysis in the R language^35^.

### 5.1 Simulation

We include a detailed description of our simulation analysis in the *Supplementary Material* of the manuscript. Here, we briefly describe that for each of ROAST, GAGE, SAFE, and CAMERA, we produce power curves for when causal transcripts are found in areas of correlation (dominating curve in Figures 1-6) or lack of correlation (lower curve in Figures 1-6). Under the same simulation scenario, we applied the JAGST competitive (Figure 7) and self-contained tests (Figure 8), and noted no power curve divergence. This indicates that there is the same power for the statistical test regardless of whether transcripts are found in correlated regions.

**Figure 1:**
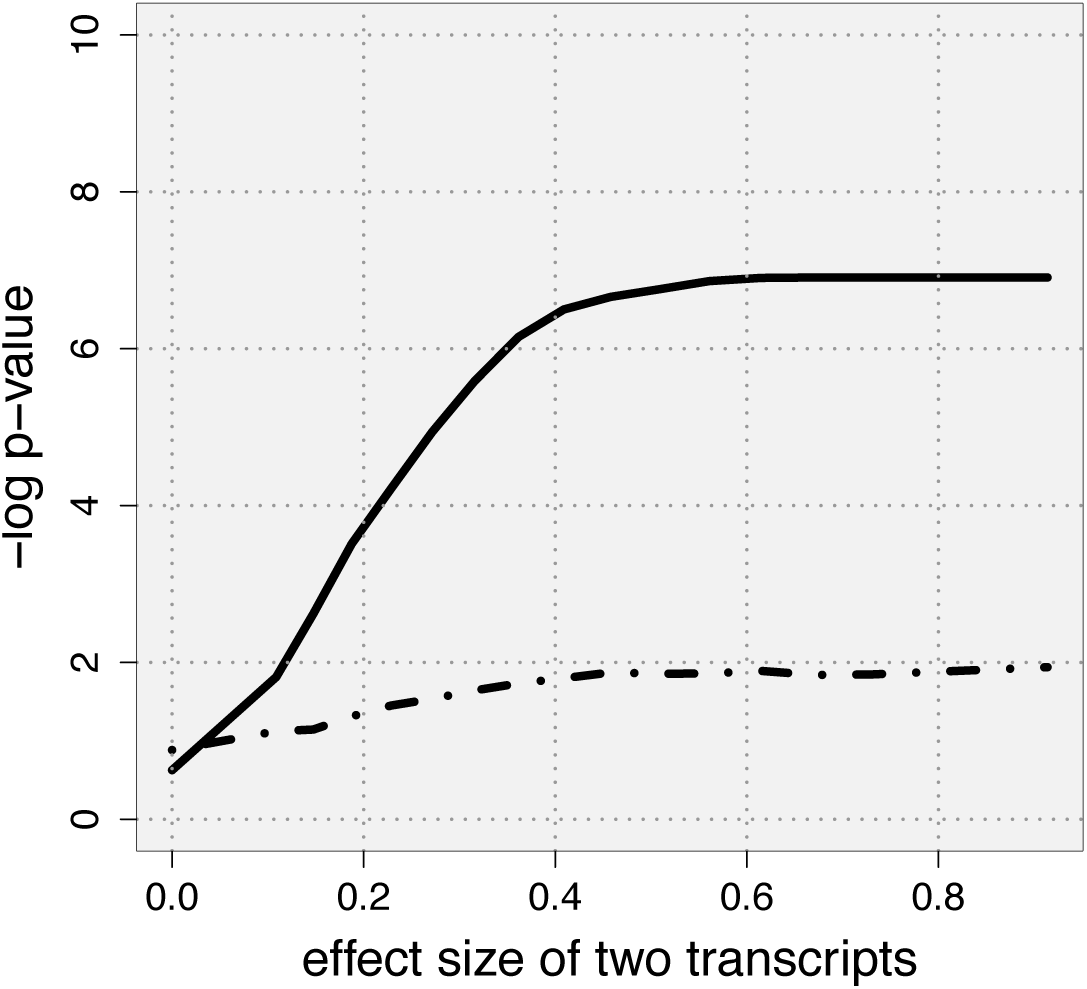
Comparison of power between SAFE tests in correlated region (solid line) and uncor-related (long dashes) region as a function of the strength of association between two transcripts within the gene set and the outcome, holding the sample size constant at 280 observations. There is a practical ceiling on the solid line curve because an empirical p-value is calculated and iterations were limited to 1000.

**Figure 2:**
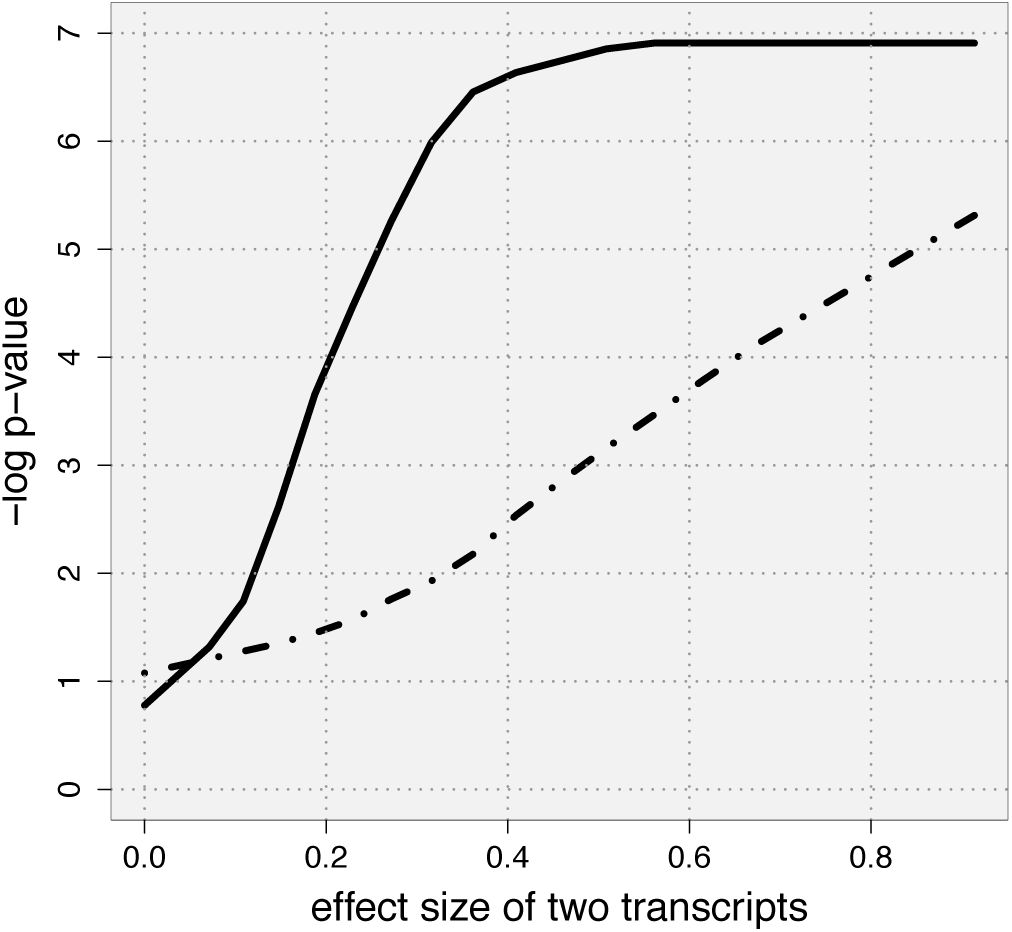
Comparison of power between ROAST tests in correlated region (solid line) and uncorrelated (long dashes) region as a function of the strength of association between two transcripts within the gene set and the outcome, holding the sample size constant at 280 observations. There is a practical ceiling on the solid line curve because an empirical p-value is calculated and iterations were limited to 1000.

**Figure 3:**
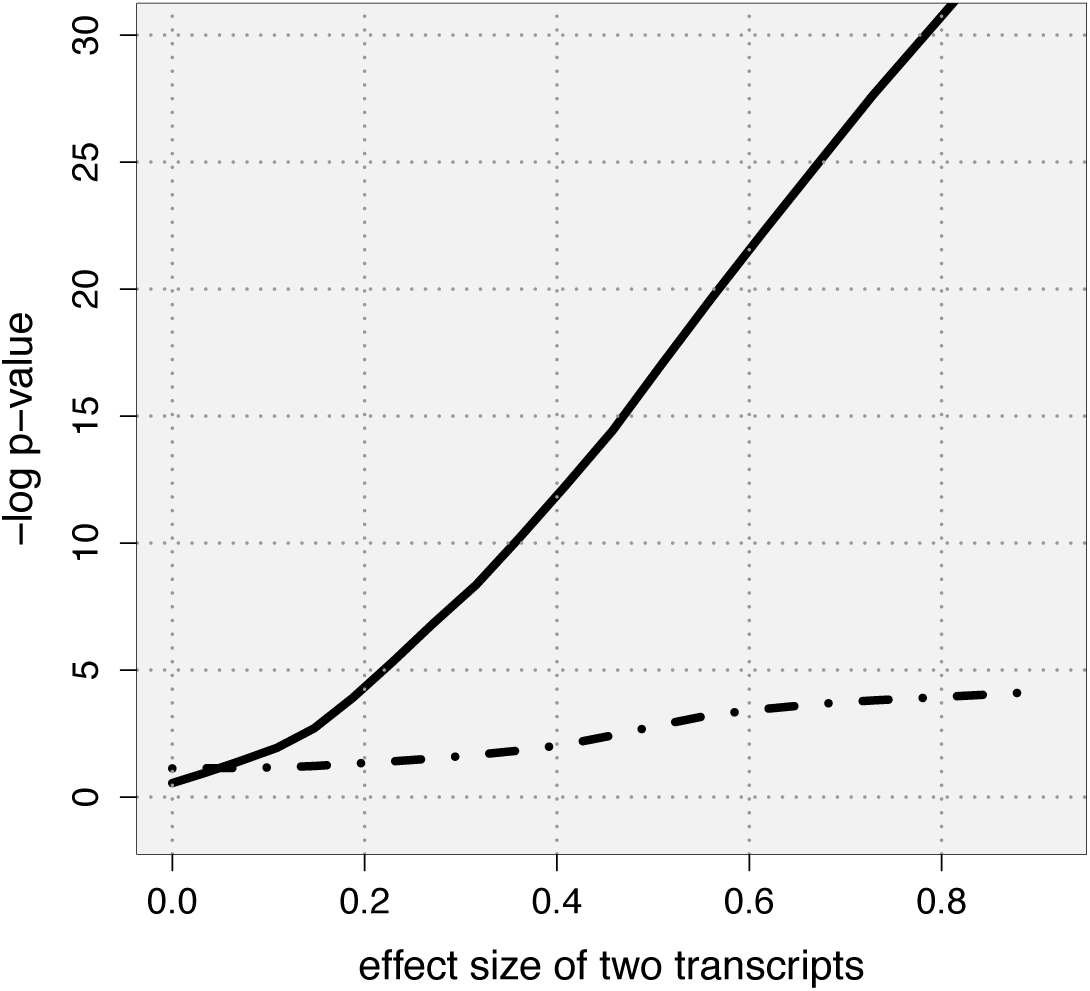
Comparison of power between CAMERA tests in correlated region (solid line) and uncorrelated (long dashes) region as a function of the strength of association between two transcripts within the gene set and the outcome, holding the sample size constant at 280 observations.

**Figure 4:**
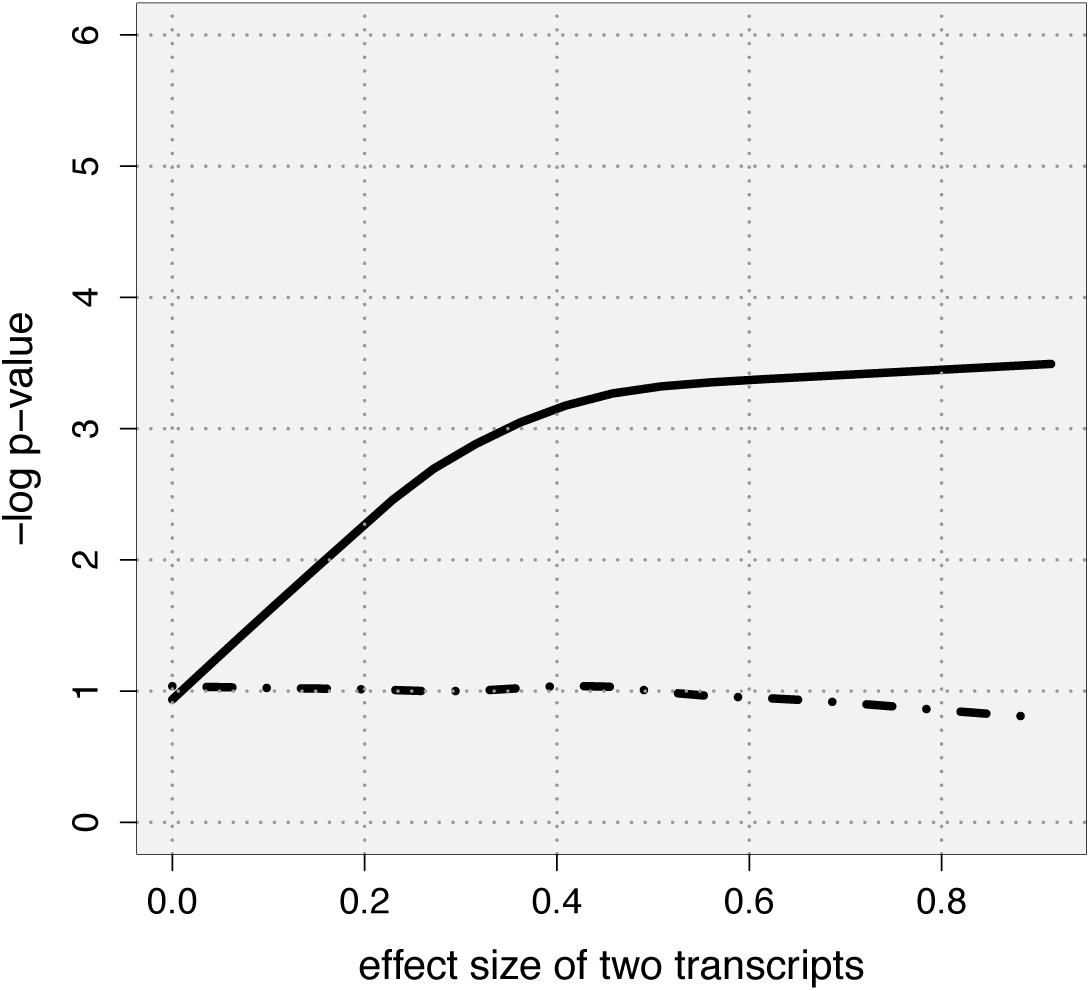
Comparison of power between CAMERA tests using ranks in correlated region (solid line) and uncorrelated (long dashes) region as a function of the strength of association between two transcripts within the gene set and the outcome, holding the sample size constant at 280 observations.

**Figure 5:**
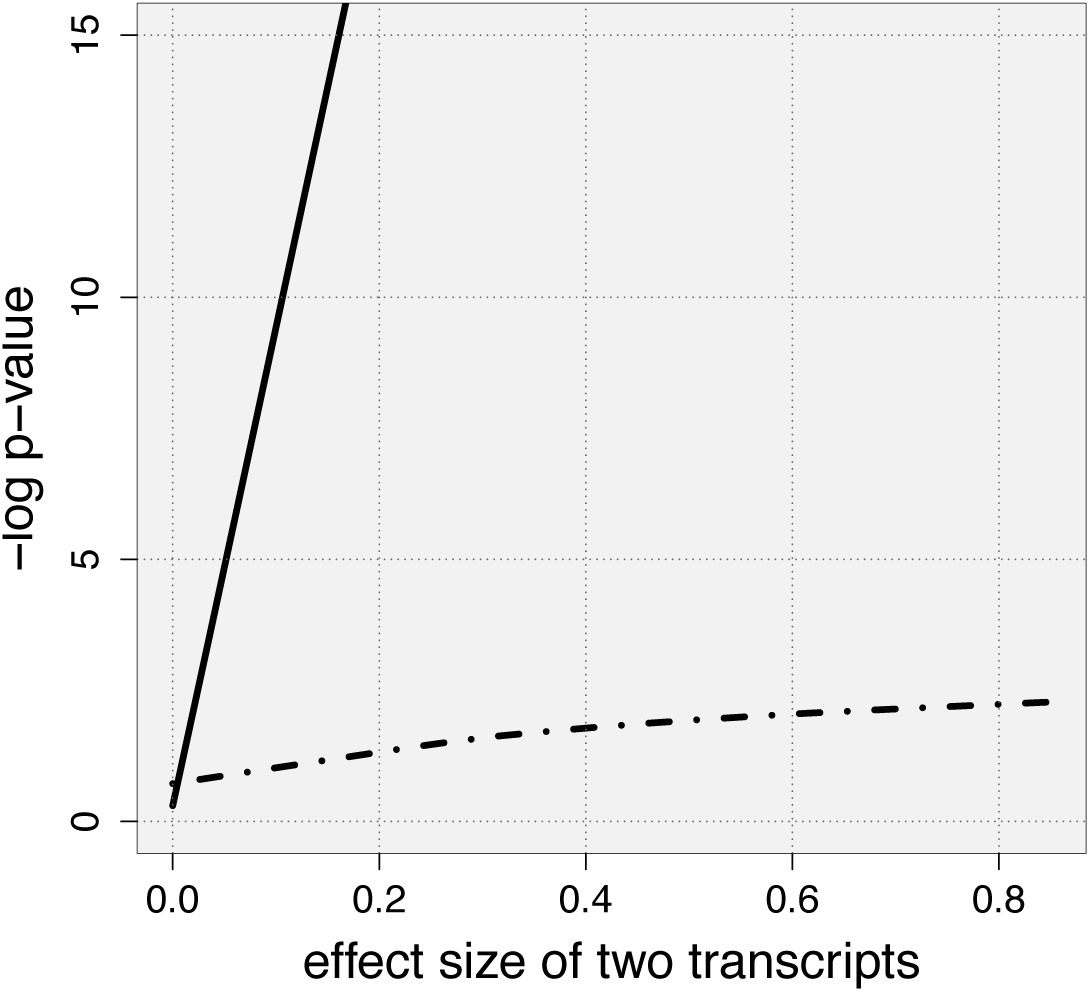
Comparison of power between GAGE tests in correlated region (solid line) and uncorrelated (long dashes) region as a function of the strength of association between two transcripts within the gene set and the outcome, holding the sample size constant at 280 observations.

**Figure 6:**
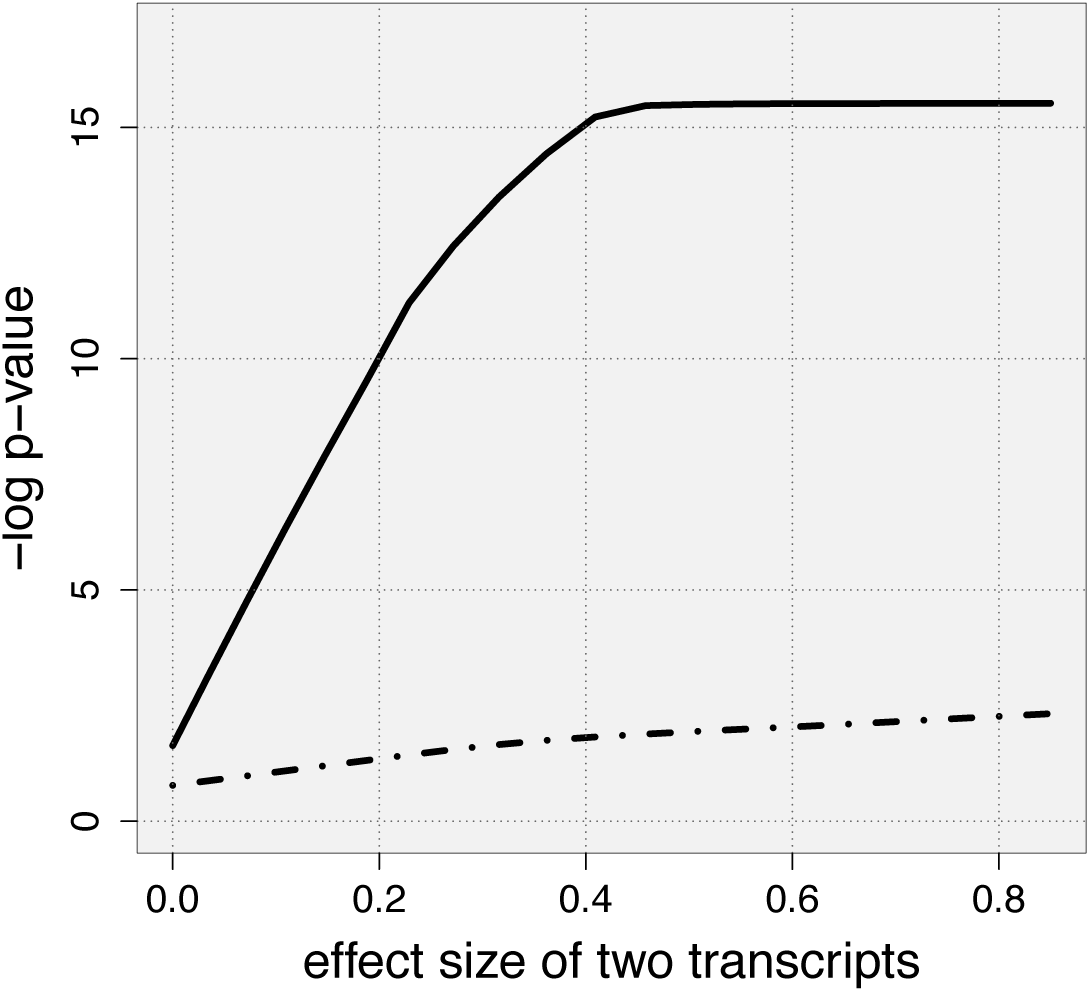
Comparison of power between GAGE tests using ranks in correlated region (solid line) and uncorrelated (long dashes) region as a function of the strength of association between two transcripts within the gene set and the outcome, holding the sample size constant at 280 observations.

**Figure 7:**
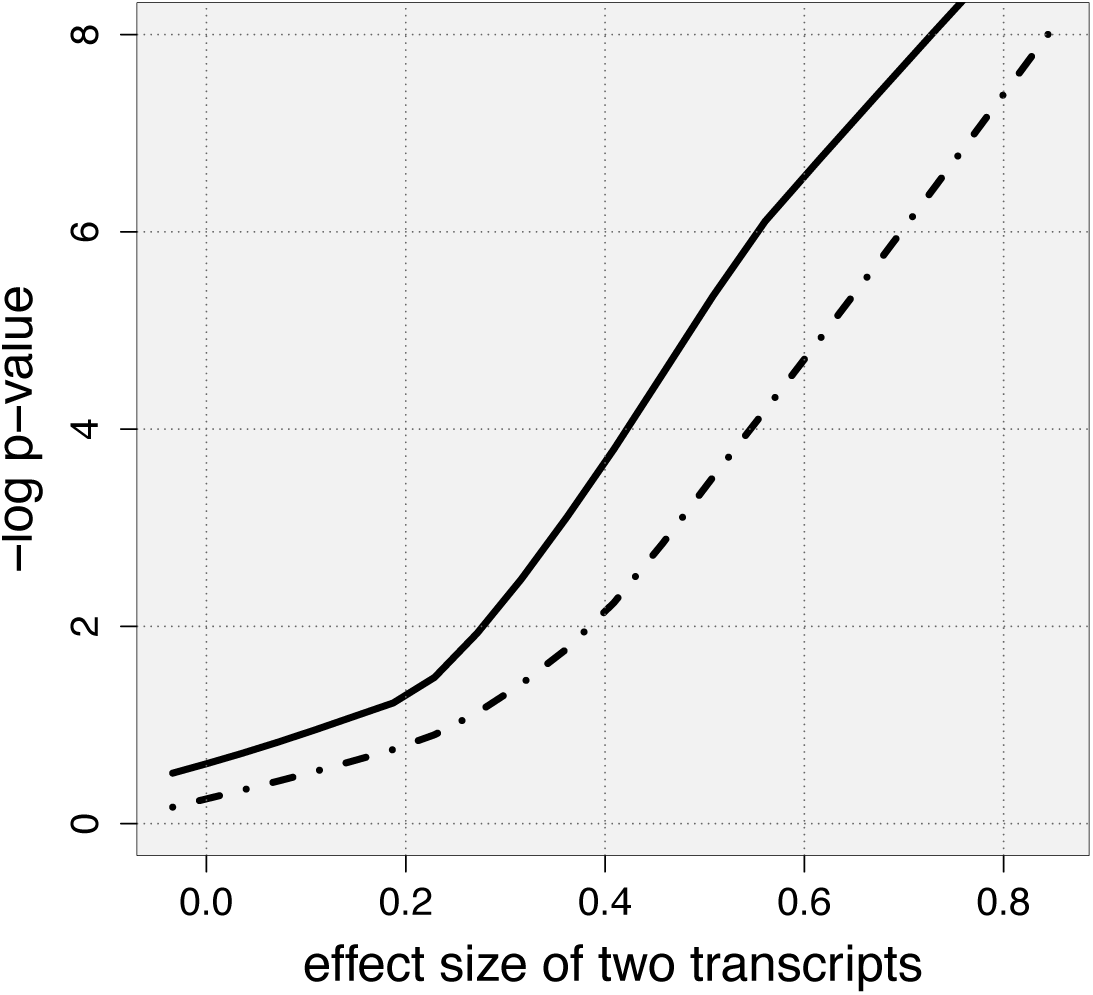
Comparison of power between the JAGST competitive test in correlated region (solid line) and uncorrelated (long dashes) region as a function of the strength of association between two transcripts within the gene set and the outcome, holding the sample size constant at 280 observations.

**Figure 8:**
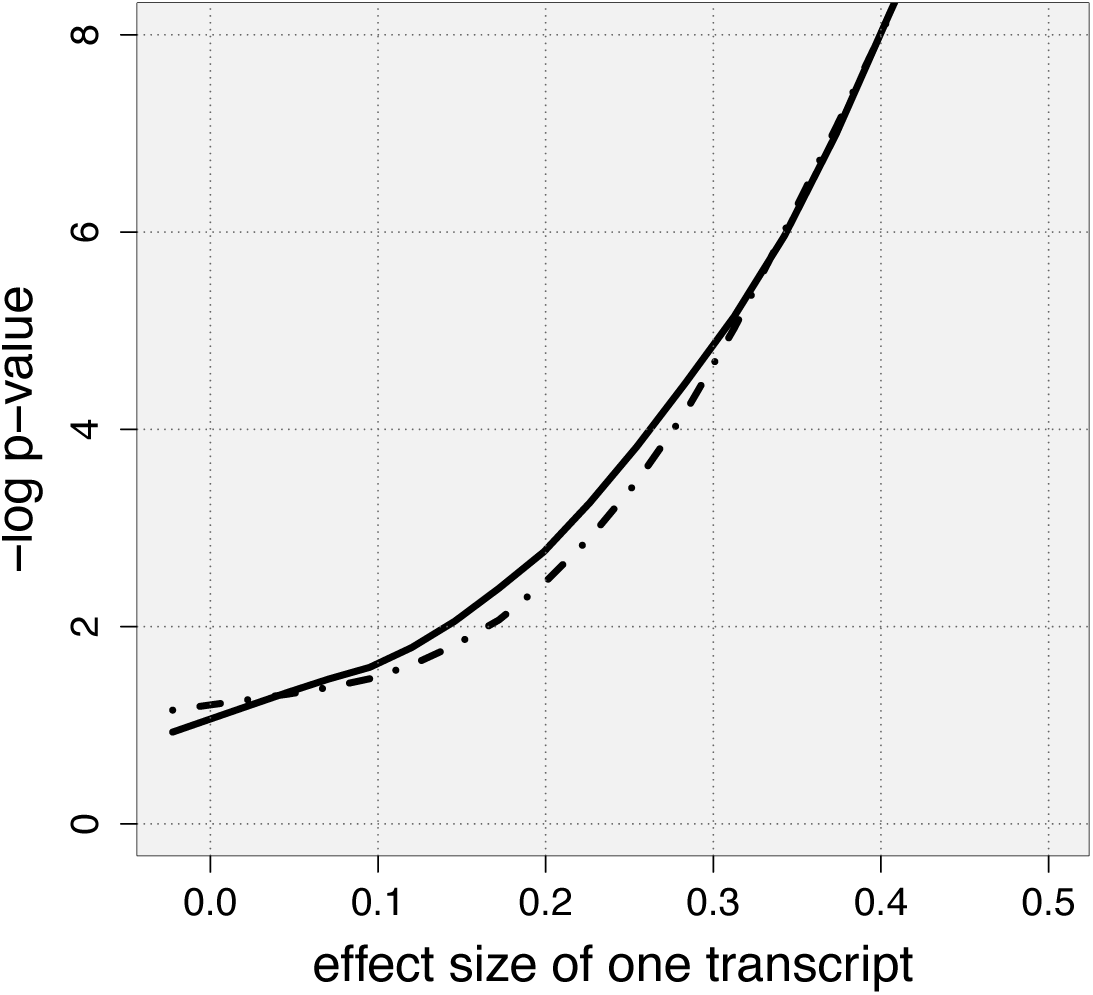
Comparison of power between the JAGST self-contained test in correlated region (solid line) and uncorrelated (long dashes) region as a function of the strength of association between one transcript within the gene set and the outcome, holding the sample size constant at 280 observations.

We also performed gene-set tests under the null hypothesis (Figure 9) and weak alternative (Figure 10) to compare results from ROAST’s directional tests with tail probabilities justified in Section 2. Comparison of resultant p-values are plotted against one another in these figures. Additional description of this simulation is given in the *Supplementary Material*.

**Figure 9:**
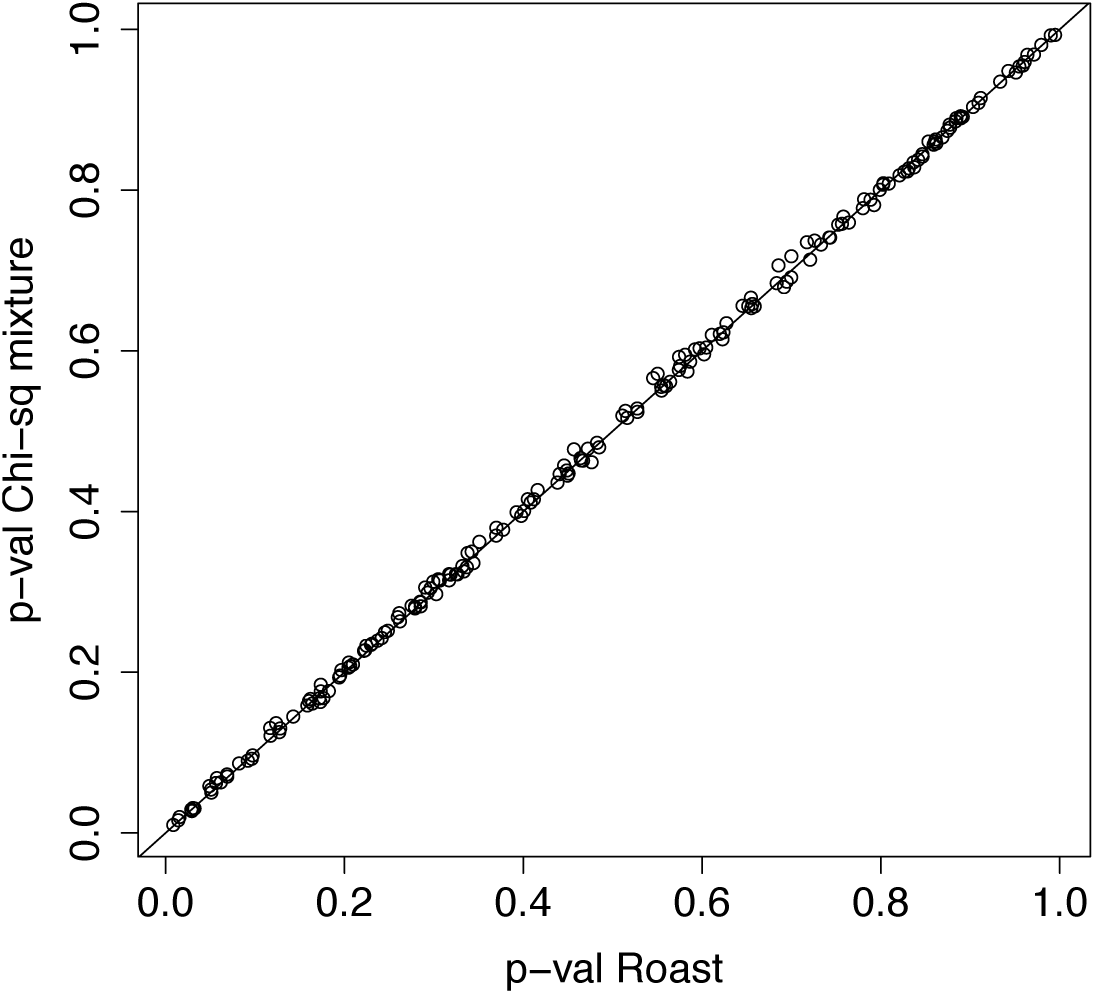
Comparison of p-values between ROAST’s directional test and tail probabilities calculated analytically. These p-values were generated under the null and have a correlation of 0.99. The line corresponds to y=x, along which equivalent points will fall.

**Figure 10:**
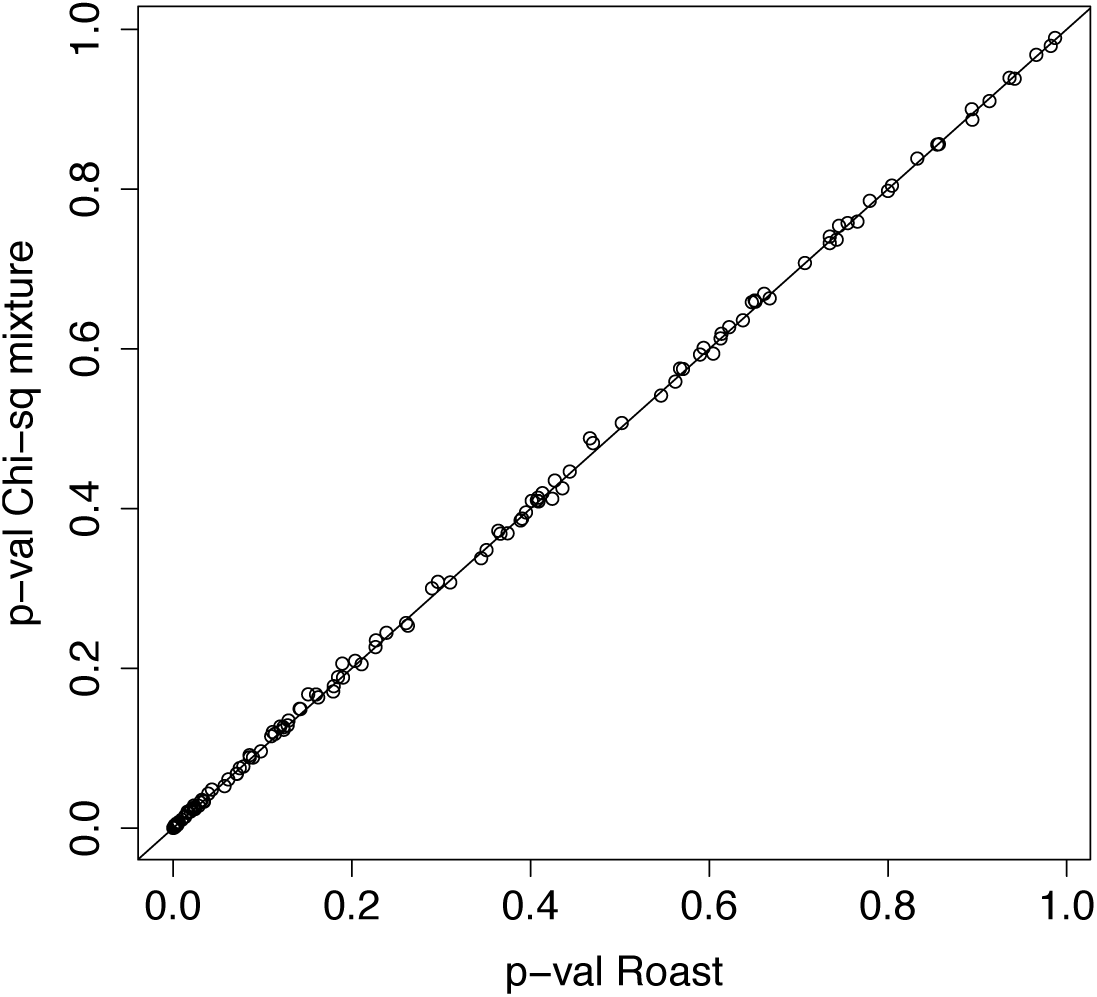
Comparison of p-values between ROAST’s directional test and tail probabilities calculated analytically. These p-values were generated under a weak alternative and have a correlation of 0.99. The line corresponds to y=x, along which equivalent points will fall.

### 5.2 Data analysis

We analyze data from an already published study of maternal and fetal transcription variation due to smoking exposure^21^. The study analyzed maternal peripheral, placenta, and neonatal cord blood on 20 pregnant women with smoking exposure and 50 without significant smoking exposure. Women were assayed using Illumina expression beadchip v3 and yielded 24,526 transcripts. We accessed the data via its Gene Expression Omnibus (GEO) accession number GDS3929^37,38^.

In keeping with the analysis protocol of Votavova et al. ^21^, we filtered transcripts of each cell type to approximately the third with a detectable level of transcript abundance. In doing so, the differential expression analysis on the remaining genes was consistent with that of ^21^ thus confirming data processing and integrity. For our gene-set analyses, we used only this approximate third most differentially expressed in each cell type category.

We analyzed 7 different gene-sets, motivated by ones commonly used in gene-set enrichment studies, oncogenic pathways, and ones more specific to the context of ^39–41,21^ (e.g., placenta development). We used 8 different gene-set test methodologies: CAMERA (with and without ranks), ROAST, SAFE, and GAGE (with and without ranks). ROAST and SAFE are self-contained tests, while CAMERA and GAGE are competitive tests. The last two used are our proposed power-invariant competitive and self-contained tests. P-values for the 8 tests by gene-set and data source (maternal peripheral blood, neonatal cord blood, or placenta) are included in Tables 1, 2, and 3, respectively. The JAGST competitive and self-contained tests are some of the least significant tests in the cases of nearly every gene-set and cell type category. This suggests that causal transcripts are correlated with many other genes in the gene-set, and our proposed tests successfully control for the correlation whereas the other gene-set tests are affected by it.

**Table 1:** P-values from different gene-set tests on maternal peripheral blood data.

| Pathway | p53 | MAPK | TGF_beta | cell_prog | plac | inflamm | immune |
| --- | --- | --- | --- | --- | --- | --- | --- |
| Camera (ranks) | 0.43 | 0.22 | 0.98 | 0.46 | 0.25 | 0.48 | 0.35 |
| Camera | 0.64 | 0.26 | 0.91 | 0.50 | 0.22 | 0.57 | 0.32 |
| Roast | 0.95 | 0.17 | 0.66 | 0.90 | 0.43 | 0.35 | 0.92 |
| Safe | 0.25 | 0.07 | 0.33 | 0.62 | 0.37 | 0.04 | 0.23 |
| Gage (ranks) | 0.14 | 0.13 | 0.34 | 0.36 | 0.28 | 0.06 | 0.12 |
| Gage | 0.08 | 0.13 | 0.22 | 0.32 | 0.42 | 0.07 | 0.12 |
| JAGST comp. test | 0.65 | 1.00 | 0.68 | 1.00 | 0.88 | 1.00 | 1.00 |
| JAGST self-cont. test | 0.21 | 1.00 | 0.26 | 0.97 | 0.58 | 1.00 | 1.00 |

**Table 2:** P-values from different gene-set tests on neonatal cord blood data.

| Pathway | p53 | MAPK | TGF_beta | cell_prog | plac | inflamm | immune |
| --- | --- | --- | --- | --- | --- | --- | --- |
| Camera (ranks) | 0.76 | 0.69 | 0.61 | 0.83 | 0.38 | 0.54 | 0.67 |
| Camera | 0.93 | 0.71 | 0.66 | 0.88 | 0.45 | 0.59 | 0.70 |
| Roast | 0.84 | 0.93 | 0.63 | 0.77 | 0.66 | 0.81 | 0.91 |
| Safe | 0.47 | 0.55 | 0.65 | 0.85 | 0.31 | 0.12 | 0.75 |
| Gage (ranks) | 0.46 | 0.58 | 0.50 | 0.92 | 0.49 | 0.17 | 0.79 |
| Gage | 0.60 | 0.57 | 0.40 | 0.85 | 0.39 | 0.11 | 0.77 |
| JAGST comp. test | 0.96 | 1.00 | 0.58 | 1.00 | 0.27 | 1.00 | 1.00 |
| JAGST self-cont. test | 0.85 | 1.00 | 0.24 | 0.99 | 0.07 | 1.00 | 1.00 |

**Table 3:**
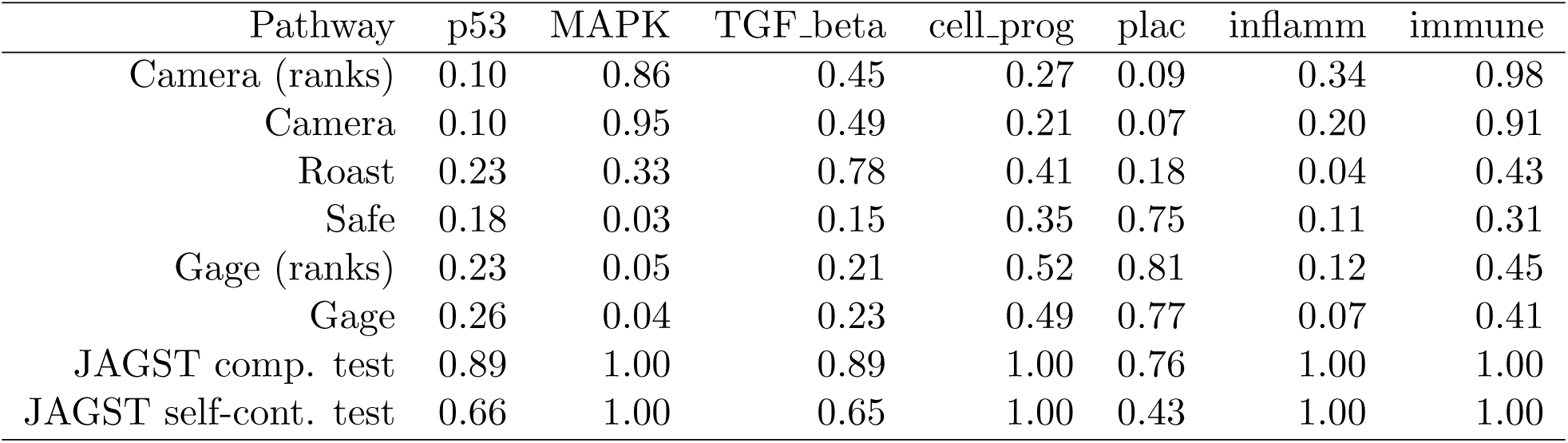
P-values from different gene-set tests on placenta data.

## 6 Discussion

Much has been learned from gene-set testing since the early 2000’s when such methods were developed. The idea that because biological processes are complex systems whose elements are not acting in isolation is powerful and has rightly been leveraged in many gene-set testing methodologies. Most of these methods are uniquely suited to certain situations, implicitly or explicitly making different assumptions of the underlying data and biological process.

We described many gene-set tests within an explicit statistical framework in this paper and showed why some could be subject to type 1 error inflation. While that observation has been made previously, its best description was done in the context of 2 x 2 tables^9^, and we describe why it can be the case with a linear combination of differential expression statistics.

More importantly in this paper, we observed and explained why all gene-set tests analyzed have variable power as a function of correlation structure. We show that if gene drivers of phenotype are found in uncorrelated regions of a gene-set, there is much less power to detect the gene-set than if the drivers are in a correlated region of the gene-set. The observation is particularly relevant because co-expression of genes on expression arrays is ubiquitous and variable; indeed, the very idea of gene-set testing is that groups of genes working in concert are expressed and can be tested together.

The gene set tests applied in our data analysis of smoking exposure on pregnant women did not yield any significant results adjusted for multiple testing. Our proposed self-contained and competitive tests were less significant in the cases of nearly every gene set, which is consistent with the idea that causal transcripts in the gene-sets tested are often correlated with non-causal ones. Since our tests control for that correlation, there is no more power for detection than if the causal transcripts were independent of non-causal ones. The proposed JAGST competitive gene set test is less significant than the JAGST self-contained test, which is consistent with the idea that competitive tests are a more significant burden of proof than self-contained tests. While we were able to replicate the differential expression results of^21^, we were not able to confirm findings of significant gene-sets. Since ^21^ relied on the DAVID database for enrichment analyses, we cannot expect our results to coincide with those of the previous study^42,43^.

The self-contained tests used in our data analysis, ROAST and SAFE, were not generally more significant than the competitive tests, Camera and Gage. Since tests are a function of correlation structure, and in the cases of the ranked tests, non-parametric, we would not necessarily expect an obvious distinction between the two kinds of tests even though their null hypotheses are different.

It is unclear why exactly there is a small divergence in power curves for the JAGST competitive test in the simulation scenarios compared (see Figure 7). Since our method relies on test statistics from an L1-penalized regression model, it is likely the relatively small variation is due to very different correlation structures from the simulation in which the transcripts with non-zero effect sizes are found. These correlation structures may affect AIC, the model fitness criterion being used to choose the tuning parameter, which then affects magnitude of the test statistics. It is important, though, that that small difference we see in our test is much smaller than that observed in other tests and that additionally the two power curves are parallel on the -log p-value scale. In contrast, we see that divergence in power curves increases with effect size for the other tests examined. For the JAGST self-contained test, we observe nearly identical power curves indicating nearly perfect power invariance.

## 7 Software and Data

An R package for implementing JAGST is available on GitHub under “sojourningNorth/JAGST” and can be installed in the standard way using directions found at the repository. Additionally, code used for simulation and analysis found in the manuscript is available at “sojourningNorth/JAGST-analysis”. Data used for analysis is available through NCBI’s GEO database and has accession number GDS3929^37,38^.

## 8 Supplementary Material

A detailed description of our simulation analysis as well as figures showing correspondences between the directional and mixed tests of ROAST and our analytical calculation of them can be found in the *Supplementary Material*.

## Acknowledgements

The author wishes to thank Prof. Arnoldo Frigessi for helpful comments in the preparation of the manuscript.

## Statistical power of gene-set enrichment analysis is a function of gene set correlation structure: Supplementary Material David Swanson

### S2 Simulation

We generated 280 samples, each with 40 transcripts or features, the 40 composed of a correlated region of size 20 with correlation 0.8 and an uncorrelated region of size 20. The correlated and uncorrelated blocks also compose our two hypothetical gene sets. We then generated two different binary outcomes consistent with a logistic regression model, where the probability of event in one case was a function of two transcripts in the correlated block and in the other case a function of two uncorrelated transcripts in the uncorrelated block. The effect sizes of the two transcripts were equal and were also the same in either scenario (i.e., whether the causal transcripts were in the correlated or uncorrelated blocks).

We varied the effect size of the two transcripts from a log odds ratio of zero to 3 in the correlated region and performed eight different gene set tests on the 20 features composing the correlated block: CAMERA (with and without ranks), GAGE (with and without ranks), ROAST, SAFE, and the two JAGST power-invariant tests were propose, one self-contained and the other competitive.

We then varied the effect sizes of the two transcripts in the uncorrelated block over the same values and performed the same eight gene tests, this time on the 20 features composing the uncorrelated block. The two power curves in each of Figures 1-6 found in the main manuscript correspond to the respective test applied to the 20 correlated or uncorrelated features and the associated outcome.

In the cases of CAMERA, ROAST, SAFE, and GAGE, at all effect sizes greater than zero, there was more power to detect the correlated block gene set, even though effect sizes were the same as compared to the uncorrelated block gene set (Figures 1-6). Indeed, the power curves diverged and had different slopes on the -log p-value scale so that there was a still bigger power difference at increasing effect sizes. The divergence is particularly striking with GAGE, whose simulation was repeated to confirm the result. In the cases of Figures 1 and 2, we see a ceiling on the -log p-value, essentially stopping the divergence of power curves. This occurs because the methods use empirical p-values, and we set the number of iterations to 1000. The divergence would continue if the number of iterations were increased. There is a similar ceiling on the rank-based tests found in Figures 4 and 6.

Our proposed gene set tests on the other hand yielded power curves that were much closer at all effect sizes of the two underlying causal transcripts (Figures 7 and 8). While there are small power differences with at least the proposed competitive JAGST test, the power curves were parallel, indicating that the power difference in curves does not increase for larger effect sizes.

Lastly we performed gene-set tests under the null hypothesis and weak alternative to compare results from ROAST’s directional tests with tail probabilities justified in Section 2 of the main manuscript. The alternative hypothesis posited two causal transcripts among a gene-set size of 20 correlated transcripts. Rather than inverting the distribution, we estimated this tail probability using the appropriate mixture distribution of central, scaled *χ*^2^’s. Figures 9 and 10 show the high degree of correspondence between the two ways of calculating the test, with a correlation of 0.99. Figures found in the Supplementary Material show the correspondence between the mixed, non-directional tests of ROAST and the analytic solution described in Section 2. Again there is a high degree of correspondence between the two tests under the null and alternative hypotheses, both with correlations above 0.9.

### S3 Figures

We show figures of the correspondence between ROAST’s directional and mixed tests and calculation of them using formulae from the Methods section of the manuscript. The alternative hypothesis assumed two causal transcripts each with log odds ratios of 0.3 and a correlated block of size 20 with correlation 0.7.

#### S3.1 Under the null hypothesis

**Figure S2:**
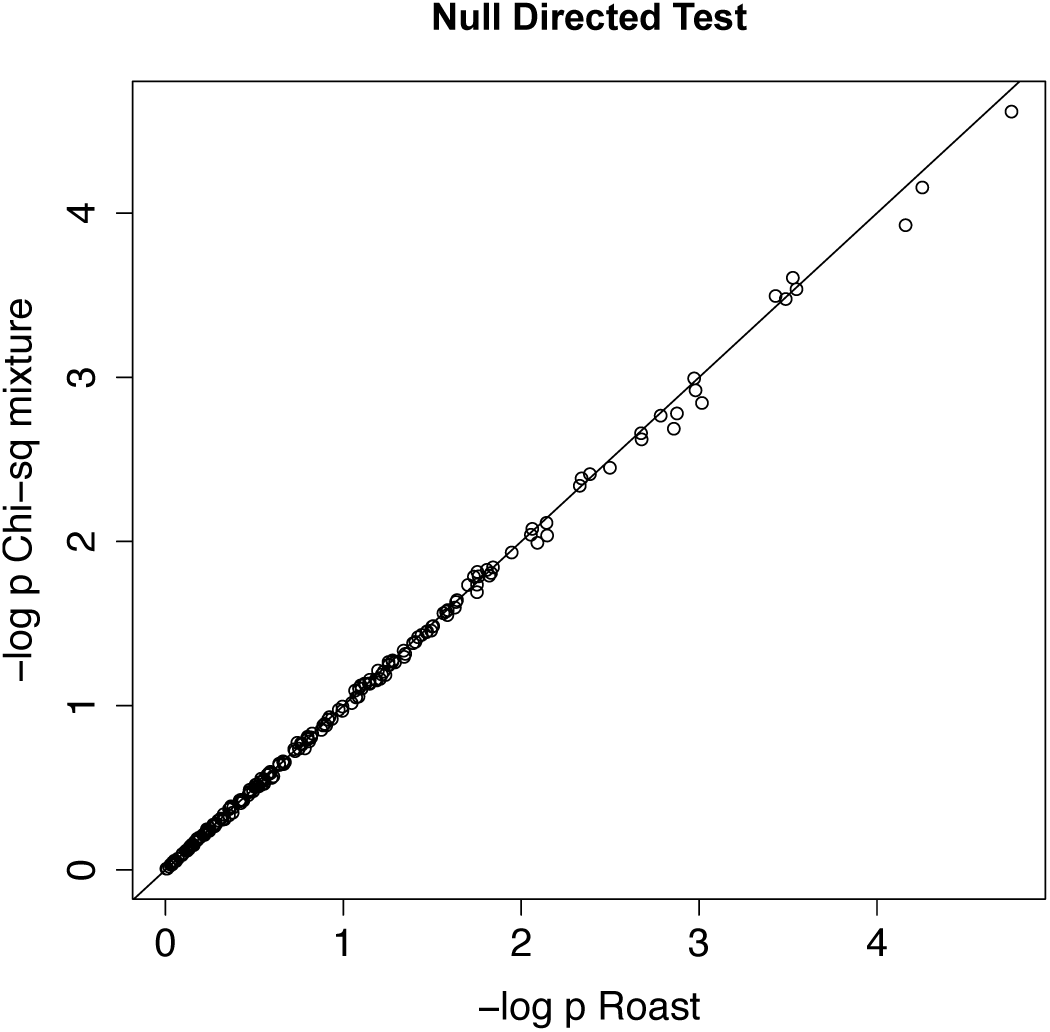
Directional test p-values under the null on the log scale.

**Figure S3:**
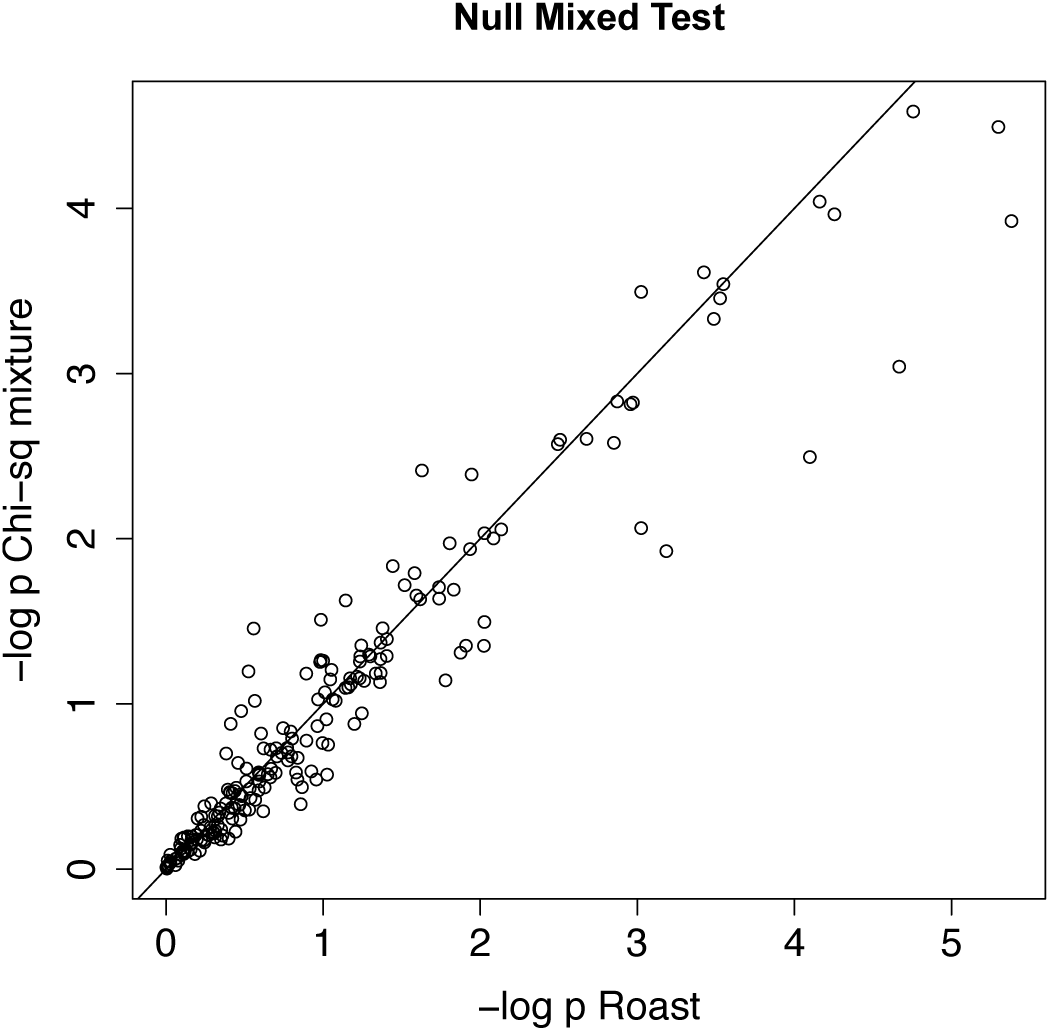
Mixed test p-values under the null on the log scale.

#### S3.2 Under the alternative hypothesis

**Figure S4:**
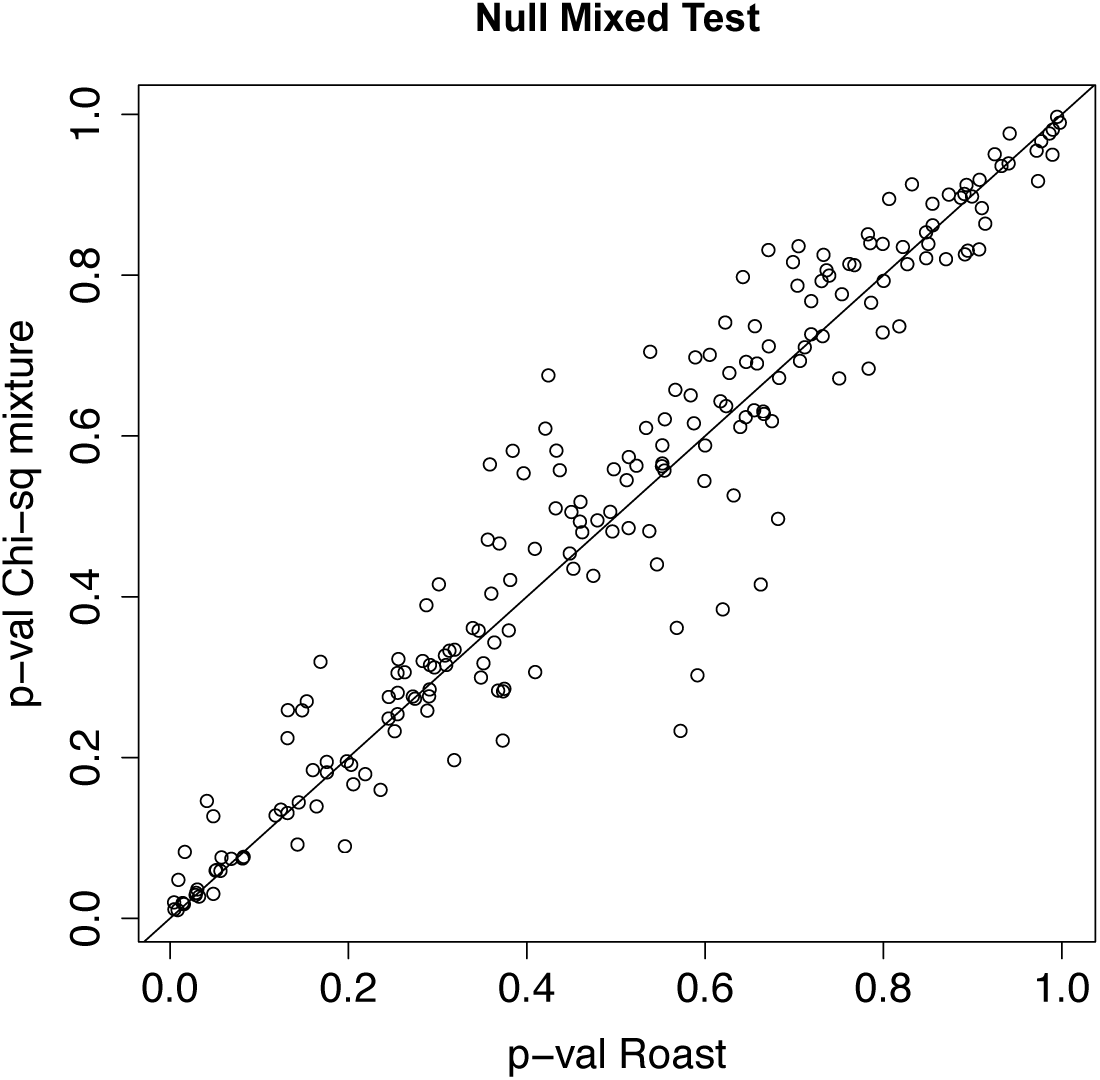
Mixed test p-values under the null on a linear scale.

**Figure S5:**
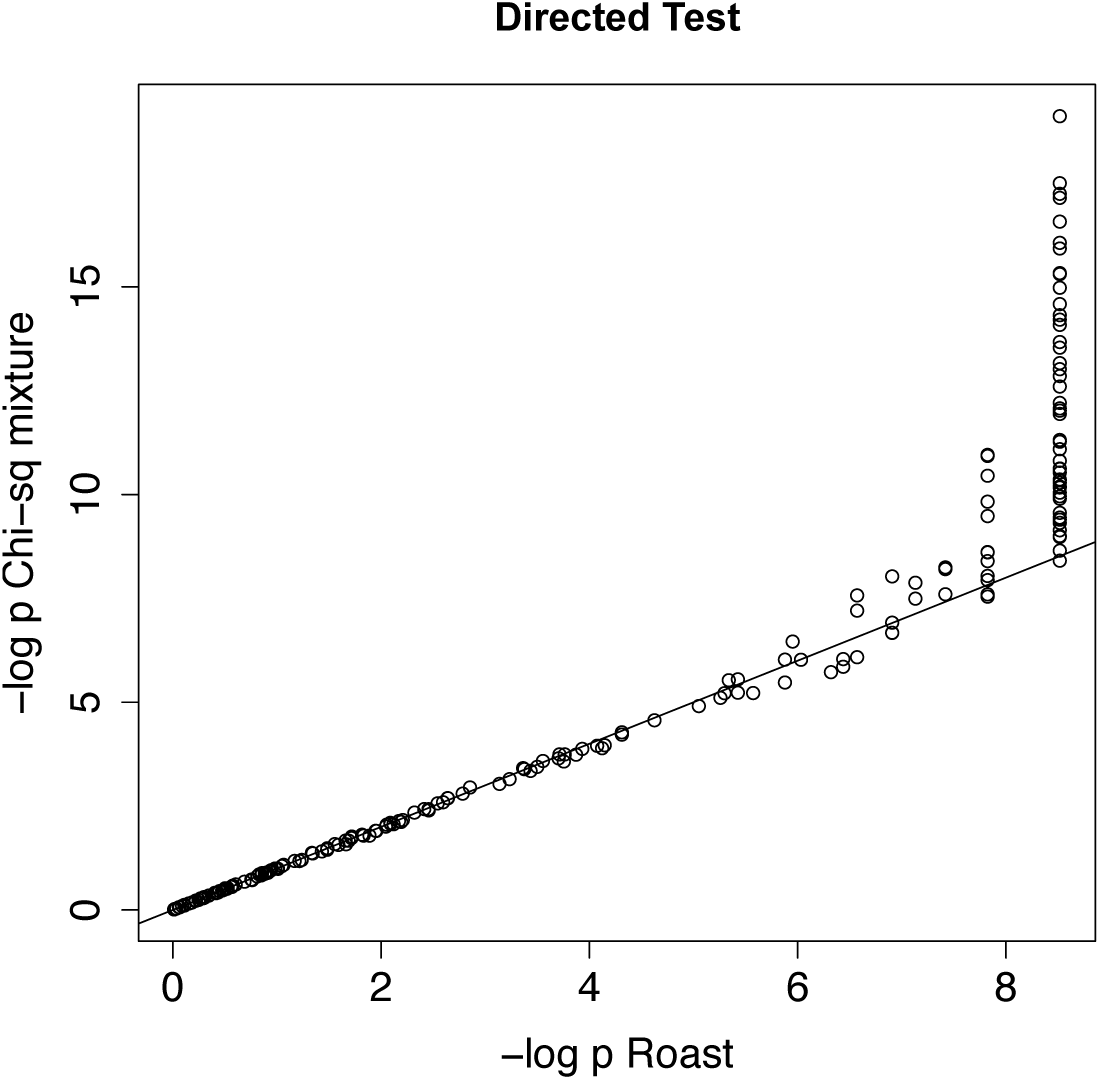
Directional test p-values under the alternative on the log scale.

**Figure S6:**
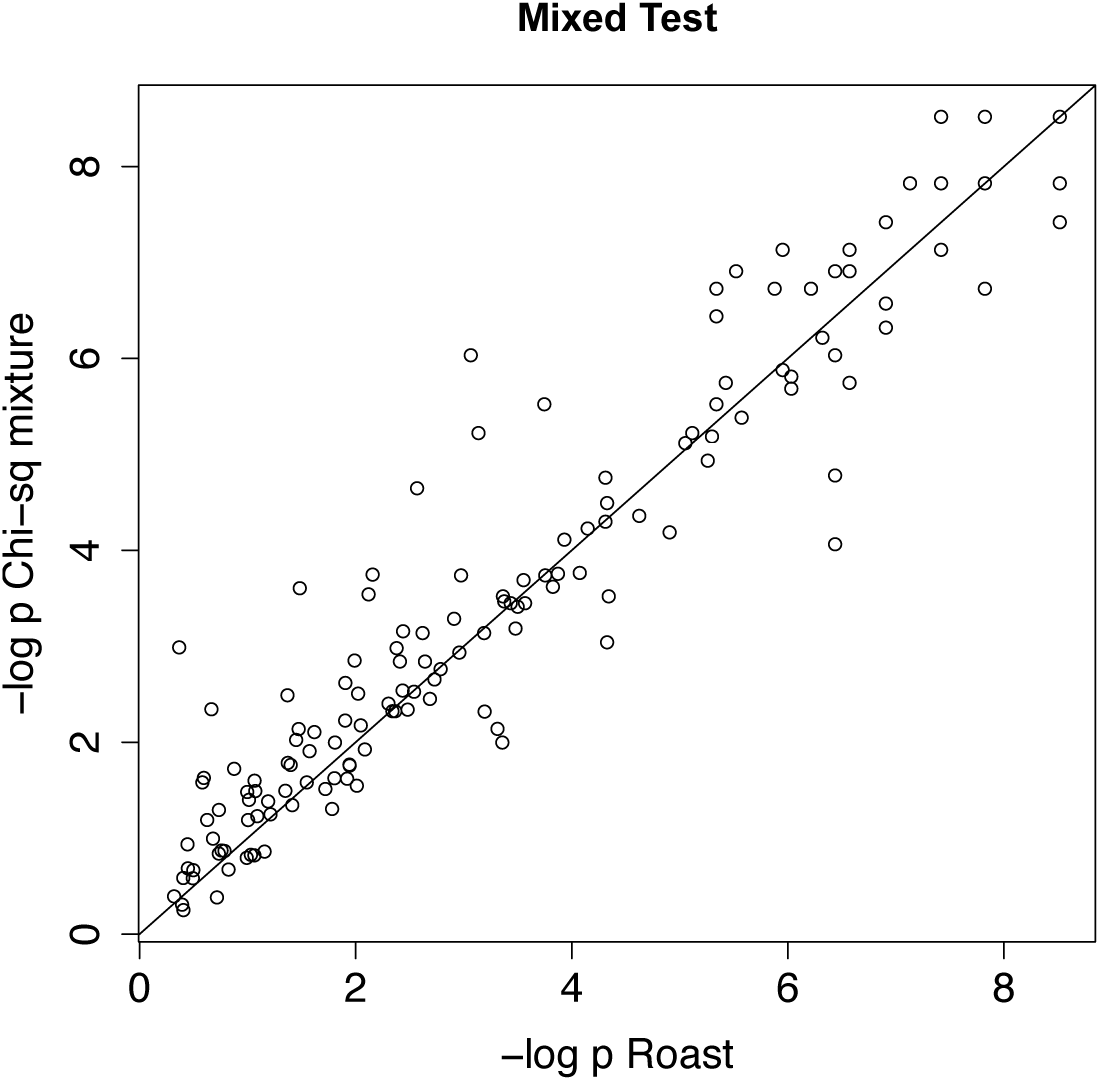
Mixed test p-values under the alternative on the log scale.

**Figure S7:**
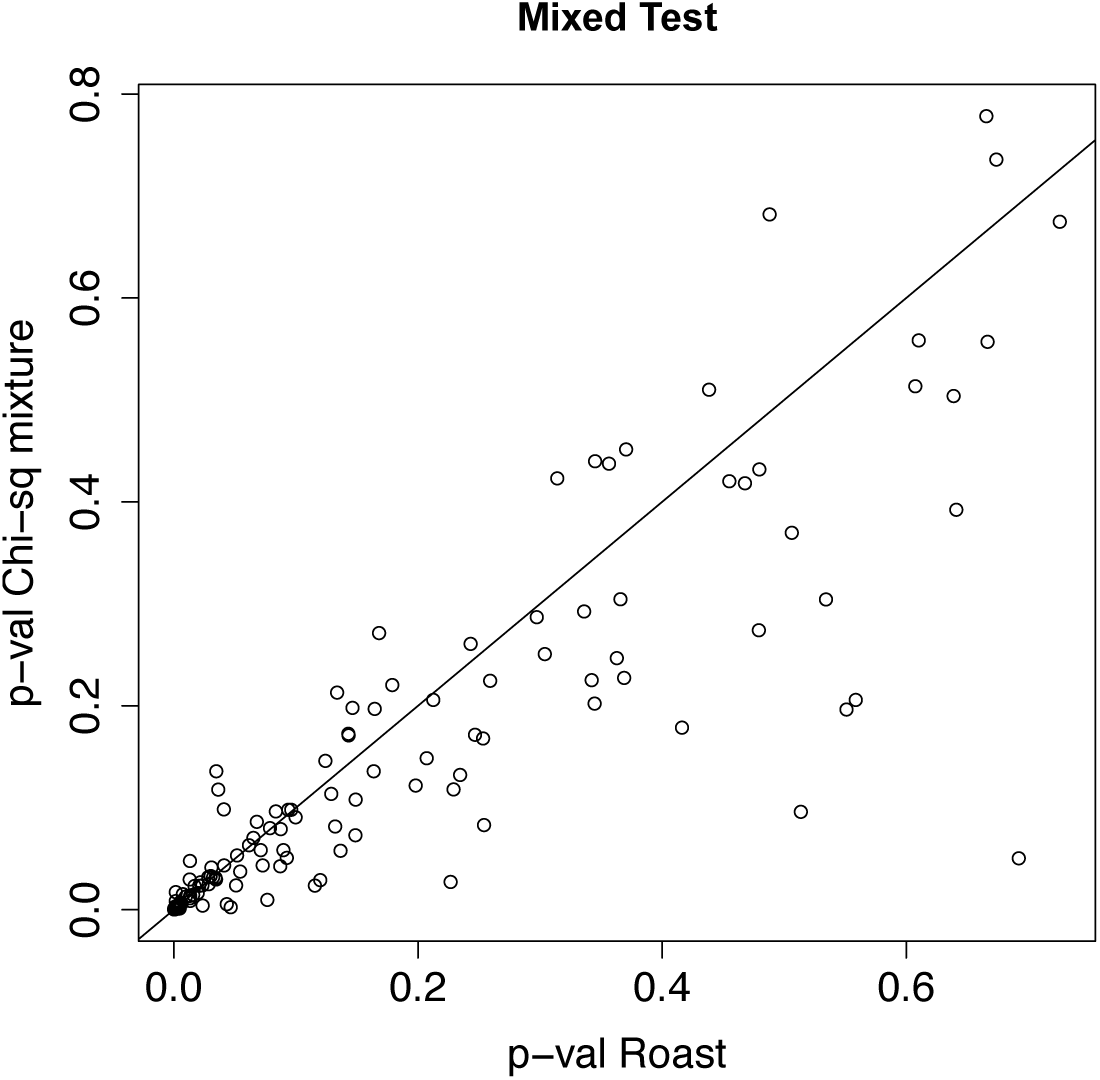
Mixed test p-values under the alternative on a linear scale.

